# A gap-filling algorithm for prediction of metabolic interactions in microbial communities

**DOI:** 10.1101/2021.05.13.443977

**Authors:** Dafni Giannari, Cleo Hanchen Ho, Radhakrishnan Mahadevan

## Abstract

The study of microbial communities and their interactions has attracted the interest of the scientific community, because of their potential for applications in biotechnology, ecology and medicine. The complexity of interspecies interactions, which are key for the macroscopic behavior of microbial communities, cannot be studied easily experimentally. For this reason, the modeling of microbial communities has begun to leverage the knowledge of established constraint-based methods, which have long been used for studying and analyzing the microbial metabolism of individual species based on genome-scale metabolic reconstructions of microorganisms. A main problem of genome-scale metabolic reconstructions is that they usually contain metabolic gaps due to genome misannotations and unknown enzyme functions. This problem is traditionally solved by using gap-filling algorithms that add biochemical reactions from external databases to the metabolic reconstruction, in order to restore model growth. However, gap-filling algorithms could evolve by taking into account metabolic interactions among species that coexist in microbial communities. In this work, a gap-filling method that resolves metabolic gaps at the community level was developed. The efficacy of the algorithm was tested by analyzing its ability to resolve metabolic gaps on a synthetic community of auxotrophic *Escherichia coli* strains. Subsequently, the algorithm was applied to resolve metabolic gaps and predict metabolic interactions in a community of *Bifidobacterium adolescentis* and *Faecalibacterium prausnitzii*, two species present in the human gut microbiota, and in an experimentally studied community of *Dehalobacter* and *Bacteroidales* species of the ACT-3 community. The community gap-filling method can facilitate the improvement of metabolic models and the identification of metabolic interactions that are difficult to identify experimentally in microbial communities.

**Author summary:** Microbes compose the most abundant form of life on our planet and they are almost never found in isolation, as they live in close association with one another and with other organisms. The metabolic capacity of individual microbial species dictates their ways of interaction with other species as well as with their environment. The metabolic interactions among microorganisms have been recognised as the driving force for the emergence of the properties of microbial communities. For this reason, understanding the effect of microbial metabolism on interspecies metabolic interactions is detrimental for the study of microbial communities. This study can be benefited by metabolic modeling and the insights offered by constraint-based methods which have been developed for interrogating metabolic models. In this paper, we present an algorithm that predicts cooperative and competitive metabolic interactions between species, while it resolves metabolic gaps in their metabolic models in a computationally efficient way. We use our community gap-filling algorithm to study microbial communities with interesting environmental and health-related applications.

## Introduction

Microorganisms form the most abundant group of living organisms on our planet. In nature microorganisms do not live in isolation, but in close association with one another, forming microbial communities. The value of microbial communities for biotechnology, the environment and human health has been profound due to their potential for use in industrial bioprocesses for the production of valuable chemicals [1], their role in biogeochemical cycles [2], and their effects on human health through the human microbiome [3].

The study of microbiomes has been extensively limited to the taxonomic classification of species and their correlation with different phenotypes. However, such correlations do not offer any mechanistic explanation on how the interactions among microbes and their environment form the observed phenotypes of the ecosystem [4]. A way to elucidate metabolic interactions in microbial communities comes from the use of a set of mathematical and computational techniques, called constraint-based methods, that make use of genome-scale metabolic models [5, 6]. Constraint-based methods have been used extensively for the study of the metabolic functions of individual microorganisms as well as for strain design in metabolic engineering [7]. In the context of microbial communities, methods like SteadyCom [8], OptCom [9], d-OptCom [10], DMMM (Dynamic Multispecies Metabolic Modeling) [11], and COMETS (Computation Of Microbial Ecosystems in Time and Space) [12] give the opportunity to evaluate growth rates and metabolic interactions of community members under various conditions.

Genome-scale metabolic models (GSMMs) can be reconstructed automatically from isolated genomes or pangenomes of specific organisms with a variety of tools [13], like ModelSEED [14] and Kbase [15], that create gene - protein - reaction (GPR) associations. However, genomes are often fragmented and contain misannotated genes [16], while the databases with information about enzyme functions and biochemical reactions used in the reconstruction process are not highly curated [13]. These problems lead to the creation of models with metabolic gaps, mass and charge imbalances and thermodynamic infeasibilities. Metabolic gaps are solved with the use of gap-filling methods, which are an indispensable part of the reconstruction process of organism-specific metabolic models [17]. The first published gap-filling algorithm was GapFill [18–20], which was formulated as a Mixed Integer Linear Programming (MILP) problem that identified dead-end metabolites and added reactions from the MetaCyc database [21] in the metabolic network. Some platforms for metabolic network reconstruction and curation, like gapseq [22] and AMMEDEUS [23], make use of more computationally efficient gap-filling algorithms formulated as Linear Programming (LP) problems. Other methods [24–26], including gapseq [22] and CarveMe [27], take into account genomic or taxonomic information in order to decide which biochemical reactions to add to the metabolic network. Other notable methods are OMNI [28] and GrowMatch [29], that try to maximize the consistency of the model with experimentally observed fluxes and growth rates respectively, while OptFill [30] is a method formulated to simultaneously solve metabolic gaps and thermodynamically infeasible cycles. Overall, most of the gap-filling methods to date are constraint-based methods that resolve metabolic gaps by adding biochemical reactions from a reference database, like ModelSEED [31], MetaCyc [21], KEGG [32], or BiGG [33], to the metabolic reconstruction of a specific organism.

Curated GSMMs of individual organisms can be combined in order to create a community model. However, GSMMs of organisms that naturally live in complex microbial communities, cannot be easily curated individually, and they often do not realistically represent the organism’s metabolic potential after the gap-filling process is completed. This problem is mainly created because of the restricted amount of physiological information that can be collected experimentally for members of complex microbial communities. Microorganisms cannot be easily cultivated individually due to their complex metabolic interdependencies with other community members. This lack of physiological information is especially acute in the case of metagenomes from the environment or from enrichments. In this paper, we propose a gap-filling algorithm that resolves metabolic gaps in microbial communities, while considering metabolic interactions in the community. The proposed community gap-filling method combines incomplete metabolic reconstructions of microorganisms that are known to coexist in microbial communities and permits them to interact metabolically during the gap-filling process. Therefore, community gap-filling can be a useful method not only for resolving metabolic gaps of GSMMs, but also for predicting non-intuitive metabolic interdependencies in microbial communities. Our algorithm also takes advantage of a formulation that ensures decreased solution times for the community gap-filling problem. Considering the unexplored metabolic potential of many natural microbial communities, as well as the difficulty to produce highly-curated metabolic reconstructions for microorganisms, we believe that this method will benefit the study of complex microbial communities.

The described community gap-filling method was applied to three independent case studies that demonstrate its ability to restore growth in metabolic models and predict both cooperative and competitive metabolic interactions between them by adding the minimum possible number of biochemical reactions from a reference database to the models. The three case studies involve a community of two *Escherichia coli* strains created for testing purposes, and the more complex communities of *Bifidobacterium adolescentis* with *Faecalibacterium prausnitzii*, and *Dehalobacter* with *Bacteroidales* species, which have interesting applications.

First, we demonstrate that our method successfully restores growth in a synthetic community comprised of two auxotrophic *Escherichia coli* strains: an obligatory glucose consumer and an obligatory acetate consumer. This community represents the well-known phenomenon of acetate cross-feeding that emerges among *E. coli* strains which grow in homogeneous environments containing glucose as the sole carbon source [34, 35].

The community gap-filling method was also used to study the codependent growth of *Bifidobacterium adolescentis* and *Faecalibacterium prausnitzii*, two well-known bacterial members of the human gut microbiome, the most diverse group of microorganisms found in the human body [36, 37]. One of the main functions of the intestinal microbiota is the anaerobic saccharolytic fermentation of complex carbohydrates such as dietary fibers [38], which leads to the production of short chain fatty acids (SCFAs) like acetate, propionate and butyrate. SCFAs are known for their various beneficial effects on the mammalian gut and metabolism [39, 40], and in particular butyrate is known for its ability to function as an energy source for the epithelial cells of the gut and works against oxidative stress and inflammation [41]. A major butyrate producer in the gut of healthy adults is the bacterium *Faecalibacterium prausnitzii* [42, 43]. The presence of *F. prausnitzii* has been connected with anti-inflammatory effects in mice [44], while its abundance is significantly decreased in some diseases of the gastrointestinal tract [45–47]. For these reasons, it is believed to have the ability to function as a probiotic and as a marker for inflammatory bowel diseases and colon cancer [48]. *F. prausnitzii* has been identified as a commensal, strictly anaerobic bacterium of the human gut that performs saccharolytic fermentation. It can utilize a variety of carbon sources from glucose, fructose, and fructo-oligosaccharides to complex molecules such as pectin and N-acetylglucosamine [49], and its typical fermentation products are butyrate, formate, lactate, and acetate [50]. *F. prausnitzii* is known for developing both competitive and syntrophic interactions with other species that live in the gut, as it competes for common carbon sources, but also has the ability to consume acetate produced by other species and convert it to butyrate. These interactions have been observed in cocultures of *F. prausnitzii* with bacterial species of the genus Bifidobacterium [51, 52]. Bifidobacterial species belong to the phylum Actinobacteria, and are commensal, obligate anaerobes of the human gut, that perform saccharolytic fermentation. They can metabolize a variety of carbohydrates [53], and their most common fermentation products are acetate, formate, lactate, succinate, and ethanol [54–56]. The cogrowth of the species *Faecalibacterium prausnitzii* and *Bifidobacterium adolescentis* has been studied both experimentally [57, 58] and computationally [59], and it has been shown that their metabolic interaction has the potential to enhance butyrate production [57]. For the community of *F. prausnitzii* and *B. adolescentis*, the community gap-filling method managed to predict metabolic interactions and SCFA production that were previously reported [57].

Finally, the community gap-filling method was used to replicate experimental results for the interaction between *Dehalobacter* sp. CF and *Bacteroidales* sp. CF50, which are the two main members of the ACT-3 community. The ACT-3 community is a microbial community found in soils and underground waters that are usually contaminated with chlorinated compounds. The study of this community has garnered scientific interest in recent years, as it has the ability to degrade chlorinated environmental pollutants [60–62]. The ACT-3 community is mainly composed of bacterial species that belong in the genera Dehalobacter, with relative abundance up to 80%, Bacteroides, and Clostridium [63]. The dechlorinating ability of the ACT-3 community mainly comes from the ability of *Dehalobacter* species to perform dehalogenation, i.e., anaerobic respiration with the use of chloroform or other chlorinated compounds as electron acceptors [64–68]. In this study, we use our metabolic reconstructions for *Dehalobacter* sp. CF and *Bacteroidales* sp. CF50, in order to simulate experimental results for the cogrowth of the two most abundant species of the ACT-3 community [63] and improve our models.

## Materials and methods

### The community gap-filling method

The general principle behind the proposed method is demonstrated in Fig 1. Our algorithm creates a compartment for each organism of the simulated microbial community, by combining the biochemical reactions of the organism’s metabolic model with those that come from a reference database. In this study, we created a curated database using the biochemical reactions available in BiGG [33]. While continuing to use the BiGG nomenclature, we removed all the biomass equations, reactions found only in eukaryotic organisms, and reactions with mass imbalances. Then, the organism compartments were allowed to exchange metabolites with one another and with their environment through a common metabolite pool where extracellular compounds can accumulate or deplete. In this way, each organism can synthesize biomass only by using its intracellular metabolites, and metabolites defined in the growth medium or produced by the other organisms of the community.

**Fig 1.**
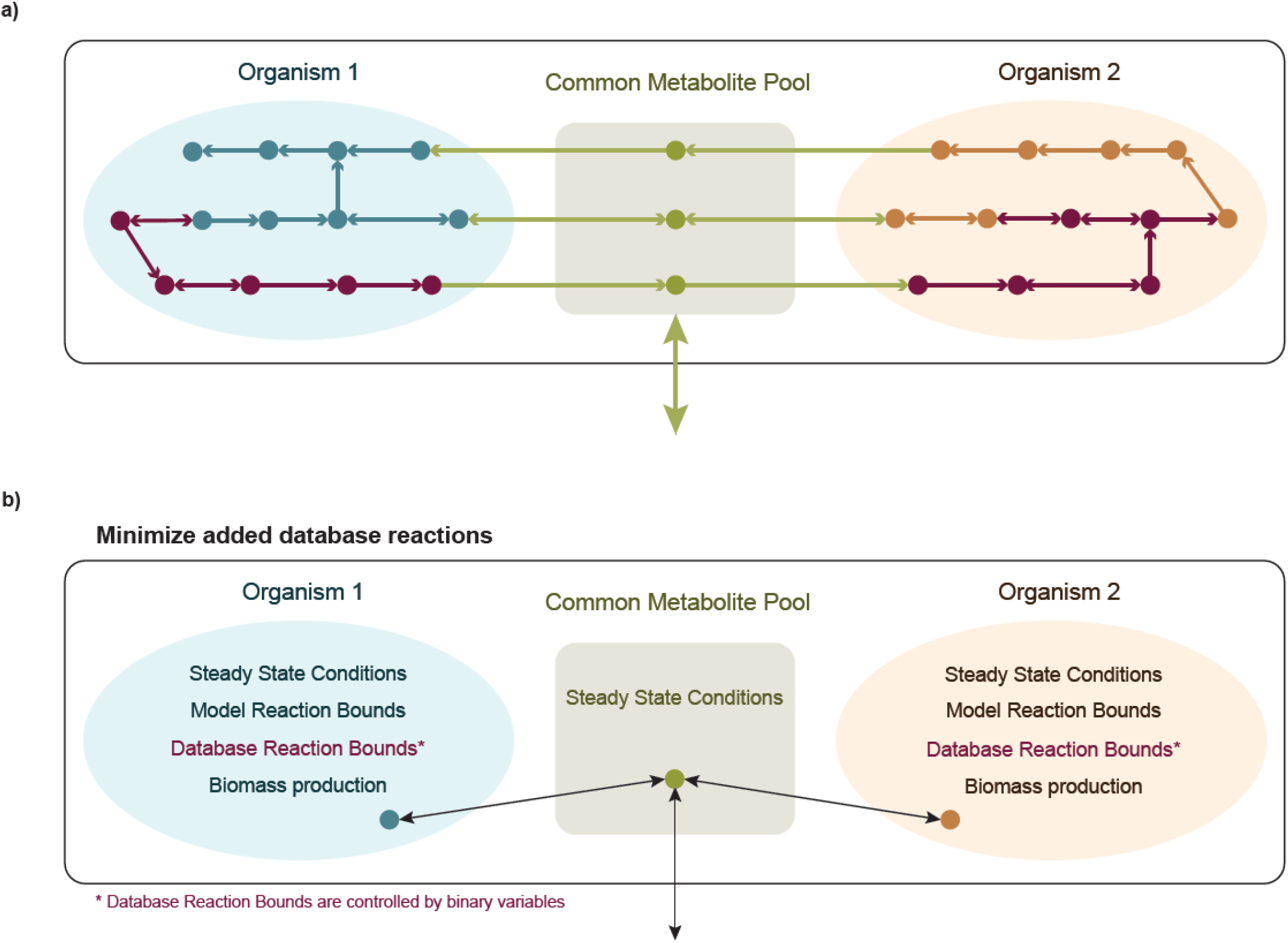
Graphical representation of the community gap-filling method. The metabolic reconstructions of two individual organisms (blue and orange respectively) are allowed to exchange metabolites and interact with the environment through a common metabolite pool (green). **a)** The algorithm adds biochemical reactions from a reference database (dark purple) to the community model. **b)** The community gap-filling algorithm is an MILP problem with the objective to add the minimum number of database reactions to the community model in order to restore biomass production in the individual metabolic models, while it satisfies some constrains for the reaction fluxes. The fate of each database reaction is controlled by a binary variable, which takes the value of 1 if the database reaction is added to the community model, and 0 otherwise.

As we can see in Fig 1, our community model consists of the organism compartments and an exchange compartment which is the common metabolite pool. This community model is used to formulate an MILP problem that predicts the minimum number of reactions that need to be added from the reference database to the community model in order to achieve a user-determined growth rate for the organisms that make up the community. The mathematical formulation of our community gap-filling algorithm for a microbial community of *N* organisms is:

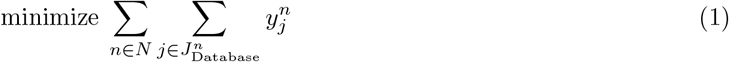

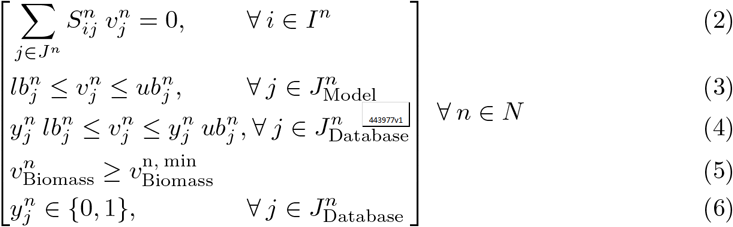

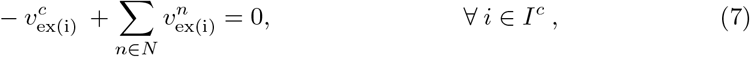

where *I*^*n*^ and *J*^*n*^ represent the number of metabolites and reactions, respectively, in the *n*^th^ microorganism compartment, while *I*^*c*^ represents the exchanged metabolites in the common metabolite pool.

Eq (1) is the objective function of the MILP problem which minimizes the total number of biochemical reactions that are added from the database to the metabolic models of the organisms composing the community. Eq (2) - Eq (7) are the constraints of the optimization problem. Specifically, Eq (2) and Eq (7) show that the metabolite pools belonging to each organism compartment and to the common exchange compartment respectively, are at steady state. Eq (3) shows the constrained fluxes of the reactions originally belonging to the organism models. The lower and upper bounds for the model reactions are selected in order to represent the growth of the organism under the desired conditions. Eq (5) represents the constraints for the fluxes through the biomass reactions of the organism models. The minimum threshold for the growth rate of each organism is defined based on existing information for the growth of the microorganism under the simulated conditions. Eq (4) implies that the fluxes of the reactions coming from the database are constrained between a lower and an upper bound, but a binary variable controls whether each database reaction will be added to the corresponding model. The binary variables, which are also used in the objective function, are defined in Eq (6). Each binary variable takes the value 1 if the database reaction is added to the community model, and the value 0 otherwise. It is noted that all the reactions from the database are initially considered reversible. Then, the minimum and maximum possible fluxes for these reactions are calculated from Flux Variability Analysis (FVA) [69] when the organism compartment is made. The FVA results are used as constraints for the database reaction fluxes, which also gives the opportunity to remove the reactions with both minimum and maximum flux zero. Using this approach, the solution space and time of the final gap-filling problem is reduced.

The simulations were performed on a machine with two 22-core Intel^®^ Xeon^®^ Gold 6238 @ 2.10GHz and 768GB of RAM, running CentOS 7. For the generation of the results presented in this paper, the MILP problem was formulated and solved in MATLAB 2017b with CPLEX Interactive Optimizer 12.8 and CobraToolbox 3, and alternative solutions for the problem were calculated with an upper time limit of two hours. The alternative solutions are used in each of the presented case studies in order to identify reactions added to the metabolic models of the participating microorganisms as well as metabolic interactions that appear more frequently in the solutions.

## Sources of metabolic models

### Core *E. coli* community

S1 Appendix explains how we used the *E. coli* core model from BiGG [70] for the creation of two *E. coli* strains, one with the ability to use glucose as a substrate and another with the ability to use only acetate excreted from the first strain. S1 Table and S2 Table contain information about reactions that were deleted from the core model and constraints for the exchange reactions that were implemented in order to create the models of the *E. coli* glucose utilizer and the *E. coli* acetate utilizer respectively. In this study, the selected lower bounds for the growth rates of the *E. coli* glucose utilizer and the *E. coli* acetate utilizer, were 0.9 and 0.09 h^-1^, respectively, based on existing information about acetate cross-feeding between *E. coli* strains [34, 71, 72].

### *Bifidobacterium adolescentis* and *Faecalibacterium prausnitzii* community

For our simulations, we downloaded the metabolic models for the strains *Bifidobacterium adolescentis* ATCC 15703 and *Faecalibacterium prausnitzii* A2-165 from the Virtual Metabolic Human Database [73]. The models are part of the AGORA version 1.03 [74], a collection of metabolic reconstructions of 773 members of the human gut microbiota. Both models are known to contain gaps in the pathways for the biosynthesis of some amino acids and vitamin B, as well as in their central carbon metabolism [73]. In this study we simulate the ability of the two models to use glucose as a substrate and exchange acetate and amino acids under anaerobic conditions. The applied constraints for the exchange reactions of the two models can be seen in S3 Table and S4 Table respectively. The lower bounds for the growth rates of the *B. adolescentis* ATCC 15703 and the *F. prausnitzii* A2-165, were set to 0.6 and 0.4 h^-1^, respectively. The constraints used for the exchange reactions of the organic acids, the amino acids, and the growth rates are based on experimental observations of *B. adolescentis* and *F. prausnitzii* strains cogrowth [57], and the study of the metabolic potential of the strain *F. prausnitzii* A2-165 [50].

### ACT-3 community

For this case study, we reconstructed the metabolic models of the two most abundant species of the ACT-3 community, *Dehalobacter* sp. CF and *Bacteroidales* sp. CF50. S2 Appendix contains information about the reconstruction process for the two models. We allowed *Dehalobacter* sp. CF to consume chloroform and *Bacteroidales* sp. CF50 to consume lactate, while the two models grow anaerobically in the same medium and are allowed to exchange organic acids and amino acids. Based on the experimental study for the coculture [63], after 18 days, the community consumes 0.5 mM of chloroform and 0.75 mM of lactate, while the total cell density is around 6·10^7^ cells·mL^-1^, and the relative abundance of the *Dehalobacter* in the community is 66% on average. This information was used in order to calculate the uptake rates of chloroform and lactate by the community as shown in S5 Table. The reaction constraints for the two models are shown in S6 Table and S7 Table. For our simulations, we used 0.1 and 0.01 d^-1^ as lower bounds for the growth rates of the *Dehalobacter* sp. CF and the *Bacteroidales* sp. CF50, respectively. All the constraints applied for the exchange reactions and the growth rates are representing experimental data from the cogrowth of the *Dehalobacter* and *Bacteroidales* species [63].

## Results

We used our community gap-filling method in order to add biochemical reactions to metabolic models of organisms that contain incomplete pathways. Since the method simulates the cogrowth of organisms that are known to coexist in microbial communities, it is used for identifying both expected and unexpected metabolic interactions in the community. In this paper, we tested the potential of the method on an artificial *E. coli* community, and then, we applied it to two communities with significance for the health of the human gut and for the bioremediation of soil and water, respectively. For each microbial community, we discuss the best solution calculated by the algorithm, i.e. the solution that adds the minimum number of biochemical reactions to the models, as well as the patterns emerging from the best alternative solutions suggested by the algorithm. The MATLAB files with all the solutions calculated by the community gap-filling for each one of the studied microbial communities can be found in the Supplementary Material. A summary of the characteristics of the used metabolic models before and after the application of the community gap-filling can be seen in Table 1. The SBML files of all the used models are available on our GitHub.

**Table 1.** Characteristics of the metabolic models used in the study before and after the application of the community gap-filling method. The algorithm simulated the aerobic growth of a community of an *E. coli* glucose utilizer and an *E. coli* acetate utilizer on glucose, the anaerobic growth of a community of the strains *B. adolescentis* ATCC 15703 and *F. prausnitzii* A2-165 on glucose, and the anaerobic growth of a community of the species *Dehalobacter* sp. CF and *Bacteroidales* sp. CF50 on media with lactate and chloroform.

| Model | Before community gap-filling |  |  | After community gap-filling |  |  |
| --- | --- | --- | --- | --- | --- | --- |
|  | metabolites | reactions | growth rate* | metabolites | reactions | growth rate* |
| *E. coli* glucose utilizer | 72 | 85 | 1.15 h<sup>-1</sup> | 72 | 86 | 0.9 h<sup>-1</sup> |
| *E. coli* acetate utilizer | 72 | 83 | 0 h<sup>-1</sup> | 72 | 85 | 0.09 h<sup>-1</sup> |
| *B. adolescentis* ATCC 15703 | 991 | 1129 | 0 h<sup>-1</sup> | 991 | 1134 | 0.6 h<sup>-1</sup> |
| *F. prausnitzii* A2-165 | 1047 | 1225 | 0 h<sup>-1</sup> | 1055 | 1233 | 0.4 h<sup>-1</sup> |
| *Dehalobacter* sp. CF | 978 | 1087 | infeasible | 978 | 1088 | 0.1 d<sup>-1</sup> |
| *Bacteroidales* sp. CF50 | 964 | 967 | 0 d<sup>-1</sup> | 965 | 969 | 0.01 d<sup>-1</sup> |

\*in the community conditions

### Core *E. coli* community

In the beginning, we tested if our community gap-filling method can restore growth in an artificial microbial community of an *E. coli* glucose utilizer and an *E. coli* acetate utilizer strain, that is allowed to grow aerobically on glucose. Our community was made after the deletion of 12 reactions from the original models of *E. coli* core metabolism (Fig 2) as discussed in S1 Appendix. As expected, the community gap-filling algorithm, instead of adding back all the knockouts that we made, suggested the addition of the minimum possible number of reactions in the community in order to restore its function. The first solution calculated by the algorithm suggested the addition of one reaction to the *E. coli* glucose utilizer model and two reactions to the *E. coli* acetate utilizer model (Fig 2). More specifically, the pyruvate oxidase (POX) reaction was added to the model of the *E. coli* glucose utilizer to produce acetate from pyruvate. The additions to the model of the *E. coli* acetate utilizer include citrate synthase (CS), which was one of the initial knockouts, and fructose-6-phosphate utilizing phosphoketolase (PKETF), which converts acetyl-phosphate to fructose-6-phosphate. Apart from the reactions that were added from the database to the models, the community gap-filling algorithm suggested the activation of reactions that were already present in the models, but did not carry flux in the FBA solution of the original core model. More specifically, in order to sustain growth in the glucose utilizer the reactions of 6-phosphogluconate dehydratase (EDD) and 2-dehydro-3-deoxy-phosphogluconate aldolase (EDA) were activated to produce pyruvate from gluconate 6-phosphate, while pyruvate dehydrogenase (PDH) was also activated to convert pyruvate to acetyl-CoA. In the acetate utilizer the preexisting reactions of acetyl-CoA synthetase (ACS) and acetate kinase (ACKr) were activated in order to convert acetate to acetyl-CoA and acetyl-phosphate respectively, and malic enzyme (ME1) converted malate to pyruvate. The fluxes for all the reactions of the community, as well as a summative table of the reactions that were added from the database to the community can be found in S8 Table. A post gap-filling FBA for the two models shows that all the biomass precursors are replenished in both models. However, both of the models grow at suboptimal rates compared to the original core model (S1 Appendix and Table 1). For the glucose utilizer, the suboptimal growth rate of 0.9 h^-1^ was due to the export of acetate that led to a reduction in resources directed towards growth, whereas for the acetate utilizer the growth rate of 0.09 h^-1^ was caused by the use of reduced acetate as the only substrate.

**Fig 2.**
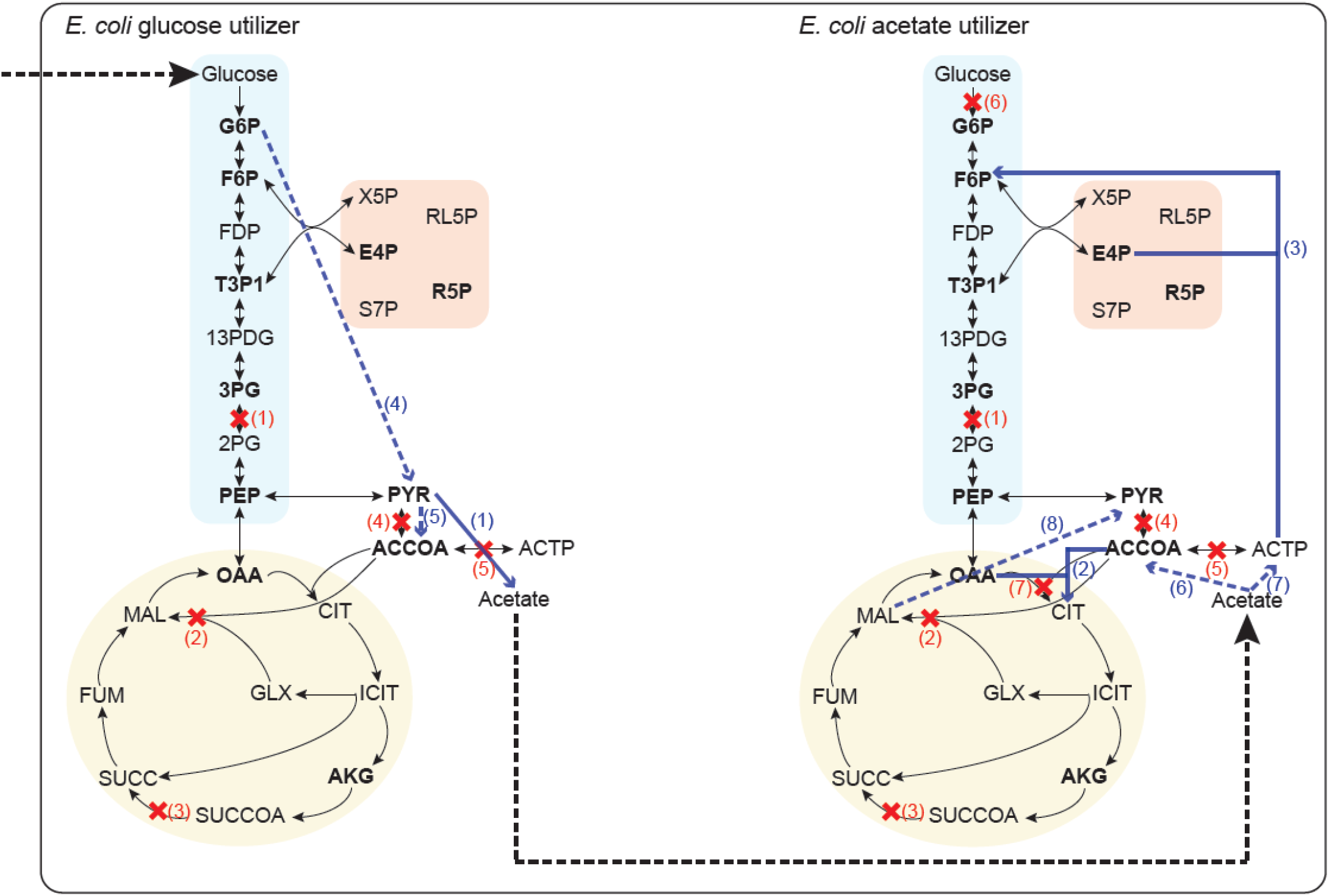
Graphical representation of the core *E. coli* community after the application of the community gap-filling method. The central carbon metabolism of an *E. coli* core model (blue rectangular: Glycolysis, pink square: Pentose Phosphate Pathway, yellow circle: TCA cycle) was used in order to create two *E. coli* strains: one that consumes glucose (left) and one that consumes acetate (right), after the deletion of the reactions marked with red crosses. The best solution of the community gap-filling algorithm predicted the addition of the reactions represented by continuous blue arrows and the activation of the existing reactions represented by dashed blue arrows, in order to restore biomass production in the two models. The dashed black arrows show the exchange reactions for glucose and acetate in the community. The metabolites in bold represent biomass precursors. The numbered deleted reactions are: (1) PGM, (2) MALS, (3) SUCOAS, (4) PFL, (5) PTAr, (6) GLCpts, (7) CS. The numbered added and activated reactions are: (1) POX, (2) CS, (3) PKETF, (4) EDD and EDA, (5) PDH, (6) ACS, (7) ACKr, (8) ME1.

The fluxes from the ten best solutions calculated by the community gap-filling algorithm for the exchange reactions of the community (S9 Table) show that all the solutions predicted the ability of the *E. coli* glucose utilizer model to uptake glucose and produce acetate that is used by the *E. coli* acetate utilizer model for aerobic or anaerobic growth. We can see that the acetate utilizing strain either shares oxygen with the glucose utilizing strain or ends up growing anaerobically due to oxygen depletion by the glucose utilizing strain in different solutions. Moreover, all the reactions that were added from the database to the community in the ten best solutions (S10 Table), carry realistic fluxes with values ranging from -10 to 15 mmol·gDW^-1^·h^-1^. We further used our artificial *E. coli* community in order to estimate the computational efficiency of our method. One major innovation of our community gap-filling method is the use of FVA before the solution of the main MILP problem. FVA is performed in order to identify the minimum and maximum fluxes that are possible for every database reaction in each organism compartment of the community. The fluxes calculated by FVA are used as constraints for the database reactions before building the community model. As a result, the solution space of the community gap-filling MILP problem is smaller and therefore, it can be solved faster. Many of the gap-filling methods to date solve MILP problems, but have not made an effort to reduce the solution space. From a computational perspective, the search of optimality can be tedious for MILP problems, and it is known that smaller feasible solution spaces can accelerate the solution of the optimization problem. Regarding the community gap-filling algorithm, the solving time increases when bigger metabolic models and databases are used and more importantly when the metabolic models of more organisms are added to the problem. As we can see from S1 Fig, the solving time of the MILP problem after the application of FVA is increasingly smaller as more organisms are added in the community, compared to the problem that was formulated immediately without the use of FVA. These results suggest that the FVA preprocessing step improves the solution time up to 84% (from 28 min to 4.5 min) for a microbial community of two organisms.

### *Bifidobacterium adolescentis* and *Faecalibacterium prausnitzii* community

In this case study, we applied the community gap-filling method to a community made from the models of the strains *Bifidobacterium adolescentis* ATCC 15703 and *Faecalibacterium prausnitzii* A2-165. The interaction between these two species has been studied experimentally and is considered beneficial since it promotes the production of the SCFA butyrate, which has antinflammatory effects in the gut [57]. Here, we simulated the anaerobic cogrowth of *B. adolescentis* and *F. prausnitzii* with glucose as the sole carbon source. None of the models can synthesize biomass in the simulated conditions, and the two models are allowed to produce SCFAs and exchange amino acids. The model of *F. prausnitzii* was permitted to either uptake or excrete acetate.

The community gap-filling algorithm restored community growth by predicting both competitive and cooperative metabolic interactions and adding database reactions to the models. The first calculated solution added five reactions to the model of *B. adolescentis* ATCC 15703 and eight reactions to the model of *F. prausnitzii* A2-165 (Table 2 and S11 Table). The reactions added to the models of *B. adolescentis* and *F. prausnitzii* participated in the biosynthesis of cofactors and amino acids necessary for growth. The first solution predicted that *B. adolescentis* and *F. prausnitzii* share the available glucose from the community medium, while *F. prausnitzii* uses part of the acetate exported from *B. adolescentis* (Fig 3). Moreover, both models export CO_2_, *B. adolescentis* exports lactate and formate in addition to acetate, and *F. prausnitzii* exports butyrate, lactate and formate (Fig 3). This correlates with experimental observations [57]. Interestingly the model of *B. adolescentis* ATCC 15703 uses the bifid shunt pathway, which is centered around the key enzyme frutose-6-phosphate erythrose-4-phosphate-lyase (F6PE4PL) in our model. The bifid shunt pathway is unique in Bifidobacteria species that commonly use it for the efficient conversion of carbohydrates, like glucose, to acetate and lactate [54, 55]. The biosynthesis of butyrate from the model of *F. prausnitzii* A2-165 happens through the reaction of butyryl-CoA:acetate CoA-transferase (BTCOAACCOAT) that converts acetate to butyrate. Beyond the expected behavior of the community regarding the use of carbon and energy sources, and the production of SCFAs, the best solution also predicts the exchange of numerous amino acids between the two community members (Fig 3). After adding the reactions suggested by the community gap-filling algorithm, FBA for the two models shows that the growth is restored, with growth rates of 0.6 h^-1^ and 0.4 h^-1^ under the simulated conditions, for the models of *B. adolescentis* and *F. prausnitzii* respectively.

**Table 2.**
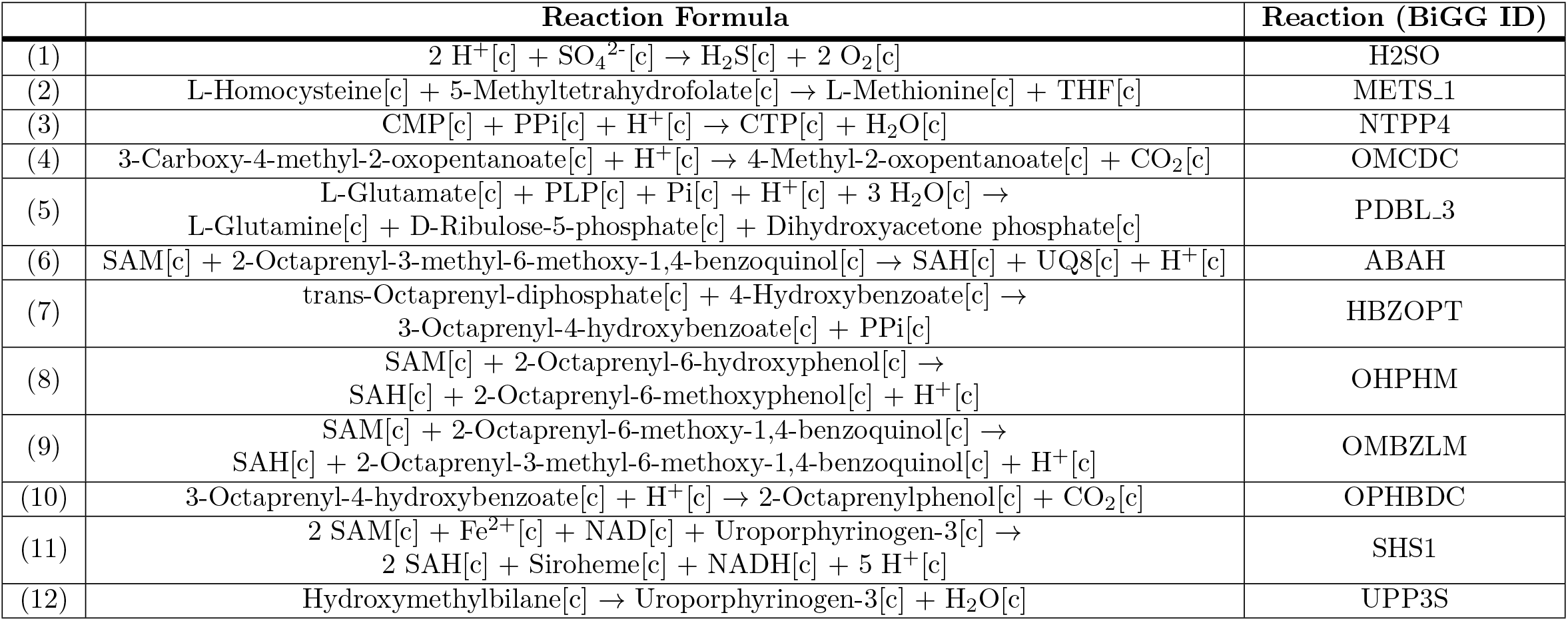
Reactions added from the database to the community model according to the best solution calculated by the community gap-filling algorithm. The reactions (1) - (5) are added to the metabolic model of *B. adolescentis* ATCC 15703, while the reactions (1) and (6) - (12) are added to *F. prausnitzii* A2-165.

**Fig 3.**
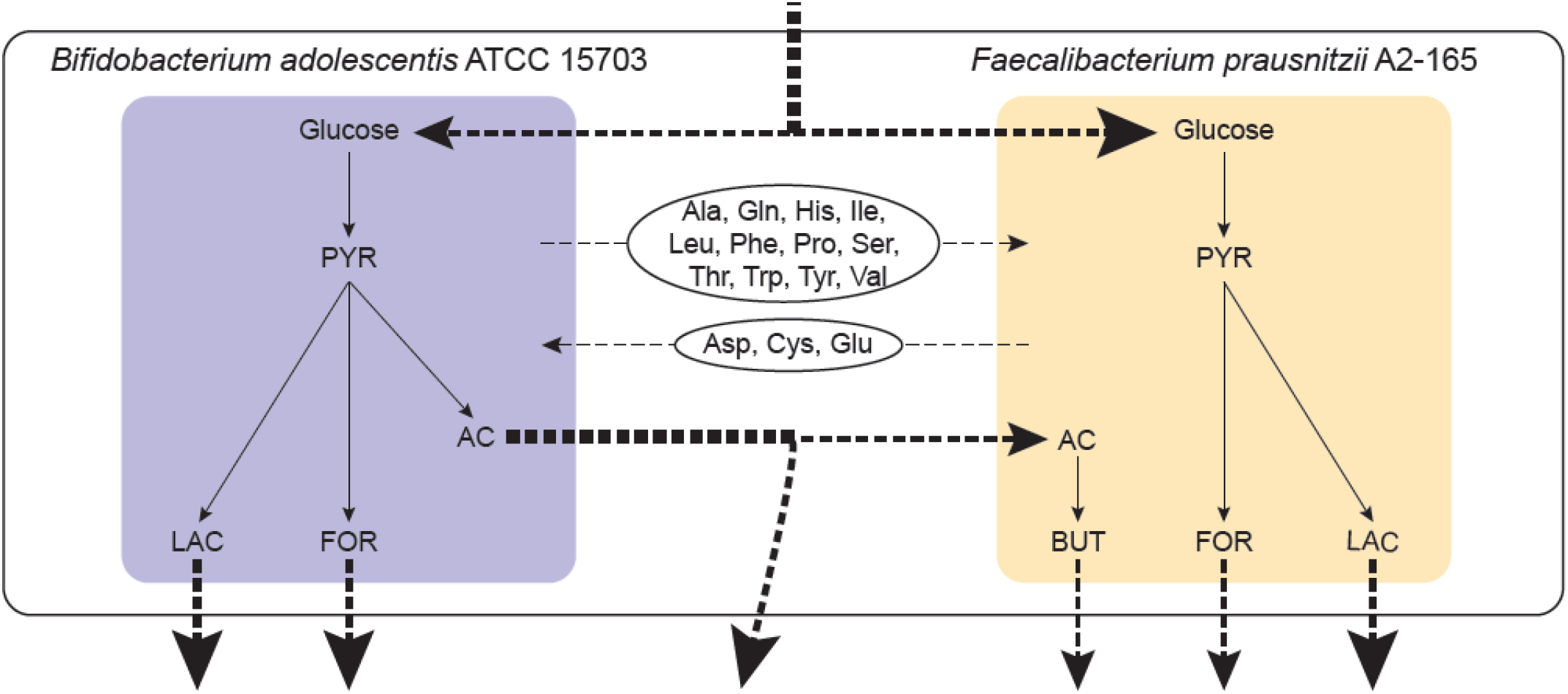
Graphical representation of the community of *B. adolescentis* ATCC 15703 and *F. prausnitzii* A2-165 after the application of the community gap-filling method. The best solution calculated by the community gap-filling algorithm predicted that the metabolic models of the strains *B. adolescentis* ATCC 15703 and *F. prausnitzii* A2-165 share the available Glucose from the common medium and produce lactate (LAC), formate (FOR), and butyrate (BUT), while they exchange acetate (AC), and amino acids (3 letter code). The non-dashed arrows represent intracellular metabolite flow and the dashed arrows represent the exchange reactions of SCFAs and amino acids. The thickness of the dashed arrows represents the relative order of magnitude of the calculated fluxes for the exchange reactions.

We also evaluated the ten best alternative solutions calculated by the community gap-filling algorithm and found several interesting interactions. First, in almost all the solutions (S12 Table), the models of *B. adolescentis* and *F. prausnitzii* shared the glucose provided in the community medium. Also, four out of ten solutions predicted that *F. prausnitzii* consumes the acetate produced by *B. adolescentis*, while the rest of the solutions predicted that *F. prausnitzii* consumes glucose from the media and produces acetate (S2 Fig). This behavior is consistent with previous studies that show the ability of *F. prausnitzii* A2-165 to function both as a consumer and a producer of acetate depending on the conditions [50]. However, to our current knowledge, acetate production from *F. prausnitzii* in the presence of *B. adolescentis* has not been observed experimentally. For this reason, more detailed research into the conditions that could affect the acetate production in cocultures of *F. prausnitzii* with *B. adolescentis* could yield interesting results.

We can see that in most of the solutions in which *F. prausnitzii* consumes acetate, it also produces CO_2_ (S12 Table, S2 Fig, Solutions 1, 3, 5), while it does not produce CO_2_ when it produces acetate (S12 Table, S2 Fig, Solutions 4, 6, 8, 9, 10). This observation is in agreement with existing information about the acetate consumption and production from *F. prausnitzii* A2-165 [50]. It is noted that in most of the solutions, CO_2_ production from *F. prausnitzii* is accompanied by CO_2_ consumption from *B. adolescentis*, while in the rest of the cases the reverse CO_2_ crossfeeding is observed. The exchange of CO_2_ between the two models could be an indication that *F. prausnitzii* and *B. adolescentis* can use CO_2_ fixation to cover their carbon needs especially in environments with limited amounts of carbohydrates. This possibility is supported by an experimental study that shows that the presence of CO_2_ in the medium promotes the anoxic growth of *B. adolescentis* species [75]. However, it is unclear if CO_2_ fixation that leads to acetogenesis can be performed through the Wood–Ljungdahl pathway in *B. adolescentis* and in *F. prausnitzii*, as it happens in other species of the gut microbiome [76, 77].

Furthermore, the production of SCFAs from the models complies with what is expected from literature [57] being that for most of the alternative solutions *B. adolescentis* produces acetate, lactate, ethanol and formate, while *F. prausnitzii* produces butyrate, fromate and succinate (S12 Table, S2 Fig). Finally, we observed patterns in the amino acid exchanges with alanine, leucine, serine and tryptophan transferred from *B. adolescentis* to *F. prausnitzii*, and aspartate and cysteine transferred from *F. prausnitzii* to *B. adolescentis* in the majority of the alternative solutions (S12 Table, S2 Fig).

Apart from the patterns that emerged regarding the exchanged metabolites in the community, the alternative solutions to the problem also indicate a consistency in the reactions that are added from the database to the model of *F. prausnitzii* A2-165 (S13 Table). However, this is not the case for *B. adolescentis*, since the reactions added from the database to the model of *B. adolescentis* ATCC 15703 are not the same through different solutions (S13 Table). Even though most of the added reactions carry realistic fluxes, some of the reactions added to *B. adolescentis* carry unrealistically high fluxes (in the order of 10^2^ mmol·gDW^-1^·h^-1^) and participate in thermodynamic infeasible cycles (S13 Table, Solutions 1 and 4, reaction PDBL 3).

### ACT-3 community

In this case study, we simulated the cogrowth of *Dehalobacter* sp. CF and *Bacteroidales* sp. CF50 that represent the most abundant bacterial species of the dehalogenating ACT-3 community [63]. Our simulation is set up in order to represent the conditions of a previous experimental study on the interaction between *Dehalobacter* and *Bacteroidales* [63]. More specifically, our community is allowed to grow anaerobically on chloroform, that can be used by the model of *Dehalobacter* sp. CF as an energy source, and lactate, that can be used by the model of *Bacteroidales* sp. CF50 as a carbon and energy source. The two models are allowed to exchange organic acids and amino acids. We show that the growth in the community of *Dehalobacter* sp. CF and *Bacteroidales* sp. CF50 can be restored by the community gap-filling algorithm.

The first solution calculated by the community gap-filling algorithm suggested the addition of one reaction to the model of *Dehalobacter* sp. CF and two reactions to the model of *Bacteroidales* sp. CF50 (Table 3 and S14 Table). More specifically, the reaction of calcium transport via the ABC system (CA2abc) was added in the model of *Dehalobacter* sp. CF. The model uses the export of Ca^2+^ through the ABC system as a proton exchange pump to generate ATP. The additions to the model of *Bacteroidales* sp. CF50 include the reactions of ammonia transport (NH4ti) and anthranilate synthase (ANS2) that are used in the model in order to import ammonia and use it to produce pyruvate from chorismate. Moreover, the first solution also predicts the ability of *Bacteroidales* sp. CF50 to uptake lactate from the medium and produce hydrogen, CO_2_, acetate, and malate that are used by *Dehalobacter* sp. CF. In parallel, *Dehalobacter* sp. CF consumes the hydrogen produced by *Bacteroidales* sp. CF50 and chloroform from the medium, and produces chlorine and dichloromethane as it performs dechlorination. In addition, *Dehalobacter* sp. CF provides pyruvate to *Bacteroidales* sp. CF50. Overall, we see that the gap-filling algorithm was able to predict the experimentally observed behavior of the community [63], and it also predicted the exchange of amino acids between the two community members, which remained elusive in the experimental study. The exchanges between the two models are depicted in Fig 4. A post gap-filling analysis shows that all biomass precursors are replenished in both models, and FBA calculated a growth rate of 0.1 d^-1^ for *Dehalobacter* sp. CF, and 0.01 d^-1^ for *Bacteroidales* sp. CF50.

**Table 3.**
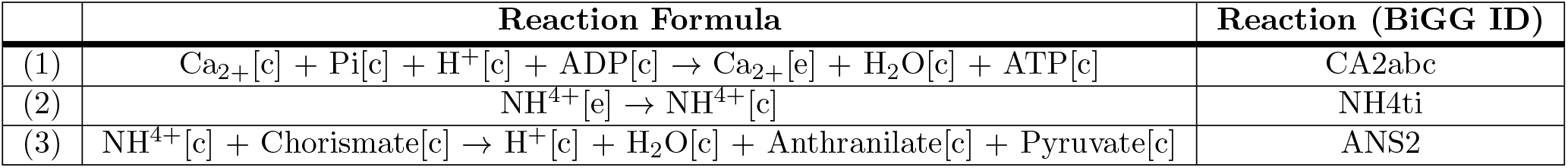
Reactions added from the database to the community model according to the best solution calculated by the community gap-filling algorithm. Reaction (1) is added to the metabolic model of *Dehalobacter* sp. CF, while reactions (2) and (3) are added to *Bacteroidales* sp. CF50.

**Fig 4.**
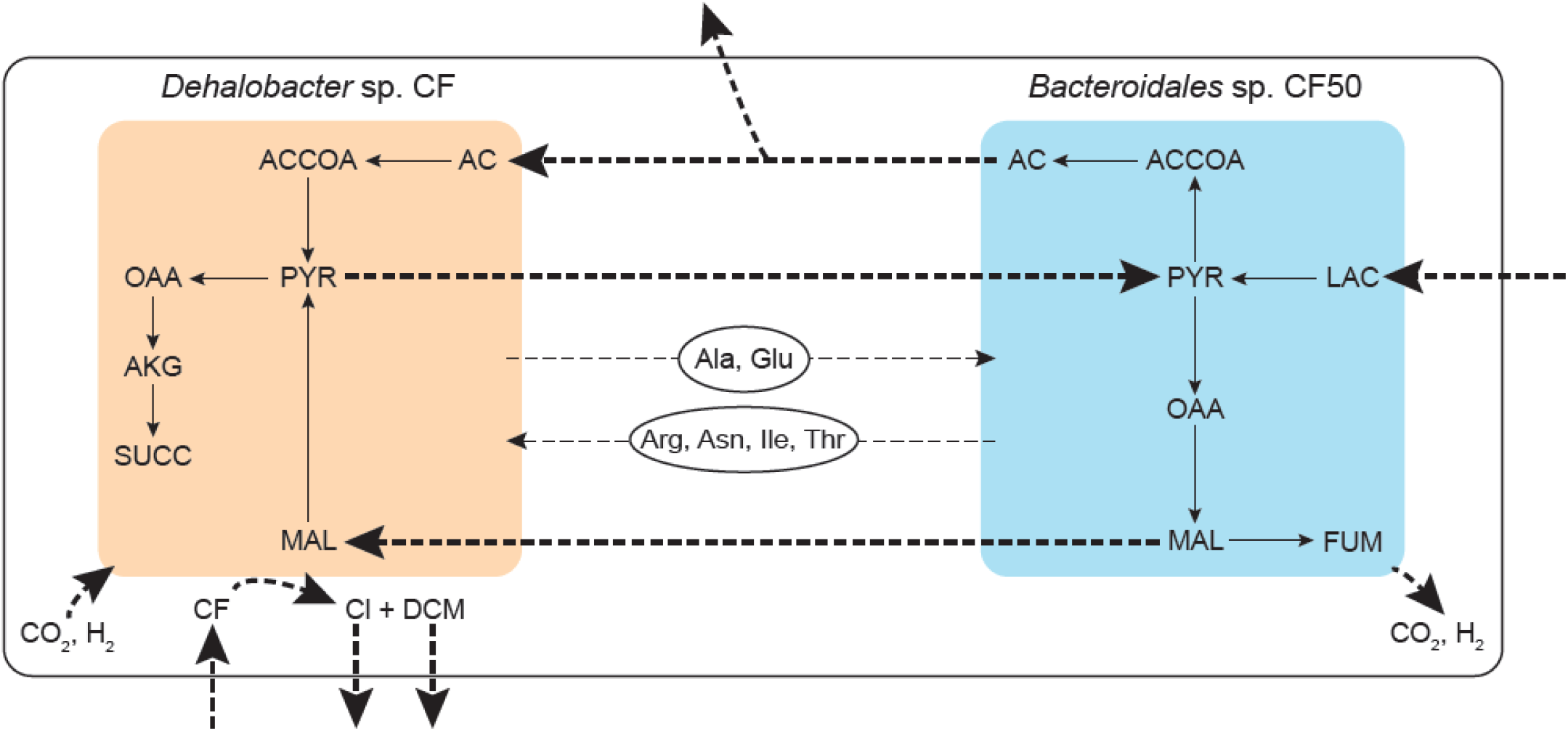
Graphical representation of the community of *Dehalobacter* sp. CF and *Bacteroidales* sp. CF50 after the application of the community gap-filling method. The best solution calculated by the community gap-filling algorithm predicted that the metabolic model of *Dehalobacter* sp. CF consumes chloroform (CF), while the metabolic model of *Bacteroidales* sp. CF50 consumes lactate (LAC) from the common medium. *Dehalobacter* sp. CF consumes H_2_ and CO_2_ produced by *Bacteroidales* sp. CF50, and uses part of the consumed H_2_ in order to respire chloroform (CF) to chlorine (Cl) and dichloromethane (DCM). The two models exchange acetate (AC), malate (MAL) and pyruvate (PYR) as expected from experimental studies. The algorithm also predicted the exchange of specific amino acids (3 letter code) between the models. The non-dashed arrows represent intracellular metabolite flow, while the dashed arrows represent exchange reactions. The thickness of the dashed arrows represents the relative order of magnitude of the calculated fluxes for the exchange reactions.

An analysis of the ten best solutions calculated by the community gap-filling algorithm shows that the model of *Bacteroidales* sp. CF50 consistently uptakes lactate, the only carbon and energy source in the medium, and ferments it into CO_2_, hydrogen, acetate, malate, fumarate and succinate, while the model of *Dehalobacter* sp. CF consistently performs dechlorination by reducing chloroform and producing dichloromethane and chlorine (S15 Table). Moreover, all the alternative solutions predicted the ability of *Dehalobacter* sp. CF to consume part of the acetate and all the malate and hydrogen produced by *Bacteroidales* sp. CF50, while providing *Bacteroidales* sp. CF50 with pyruvate (S15 Table, S3 Fig).

In most of the solutions *Dehalobacter* sp. CF consumes part of the CO_2_ produced by *Bacteroidales* sp. CF50 (S15 Table, S3 Fig, Solutions 1 - 5, 8, 10). This exchange could indicate that *Dehalobacter* sp. CF performs CO_2_ fixation, which is possible since the species is known to possess the Wood–Ljungdahl pathway [63]. Furthermore, we observed patterns in the amino acid exchanges between the two community members, with arginine, asparagine, isoleucine, lysine, phenylalanine, threonine, and tyrosine transferred from *Bacteroidales* sp. CF50 to *Dehalobacter* sp. CF, and alanine and glutamate transferred from *Dehalobacter* sp. CF to *Bacteroidales* sp. CF50 in the majority of the alternative solutions (S15 Table, S3 Fig).

More information about the quality of the solutions calculated by the community gap-filling algorithm can be retrieved by closer inspection of the reactions that were added from the database to the models in the ten best solutions. We can see that in most of the solutions (S16 Table, Solutions 1 - 6, 8, 10), even though the reactions added to the model of *Bacteroidales* sp. CF50 have realistic fluxes, those added to the model of *Dehalobacter* sp. CF carry unrealistically high fluxes (close to the infinite reaction bounds). For example, the already discussed reaction of calcium transport via the ABC system (CA2abc), which is used for ATP production in the model, has unrealistically high fluxes and forms a cycle with the reaction calcium transport in/out via proton antiporter (CAt4) that already exists in the *Dehalobacter* sp. CF model (S16 Table, Solutions 1, 2, 3).

## Discussion

Our results demonstrate the ability of the community gap-filling method to resolve metabolic gaps and restore growth in metabolic models by adding the minimum possible number of biochemical reactions from a reference database to the models and activating already existing reactions of the models, while the models are allowed to interact metabolically. In the first test case study, the algorithm restored community growth and predicted the exchange of acetate between the two *E. coli* strains. For the community of *B. adolescentis* and *F. prausnitzii* in the second case study, the algorithm calculated the competitive consumption of glucose present in the common medium, the syntrophic exchange of acetate, CO_2_ and amino acids, and the production of SCFAs. The algorithm also identified the exchange of organic acids, amino acids, CO_2_ and H_2_ for the community of *Dehalobacter* and *Bacteroidales*. In all three case studies, the metabolic exchanges were not strictly forced with constraints, but they emerged from the algorithm. We also showed that the implementation of FVA in our algorithm resulted in significantly decreased solution time of the MILP community gap-filling problem.

In general, the community gap-filling method could facilitate the study of complex communities where the existing information for the involved species is incomplete. A method like this can be used at the last steps of the metabolic reconstruction process, and it can also become part of the efforts for the creation of a microbiome modeling toolbox [78]. However, like every other constraint-based method, our community gap-filling method has some weaknesses. One of the most prominent challenges is that the performance of the algorithm depends highly on the quality of the input models and databases being used. More specifically, the absence of some important reactions from the models can lead to poor predictive power, while unconstrained reaction directionality, especially for exchange reactions, can affect the predicted metabolic interactions and give birth to flux-carrying cycles between the community members.

The algorithm performs better with highly curated models, but even then it shows preference over specific pathways. This bias comes from the fact that the process of manual curation and gap-filling for a model is based on the use of specific media and requires knowledge about the growth of the organism under specific conditions. In addition, the composition and the quality of the database reactions is important. Ideally the database should contain balanced biochemical reactions that represent a diversity of metabolic functions observed in prokaryotic organisms, or any type of organisms that represents the models being used. Regarding the solutions of the algorithm that predict reactions with unrealistically big fluxes and reactions that form thermodynamically infeasible cycles, they can either be excluded completely from the analysis of results as they are not trustworthy, or they could be avoided by further constraining the reactions of the metabolic models based on thermodynamic information [79, 80]. From the perspective of computational efficiency, the total running time of the algorithm can be further decreased with the use of newer and faster parallel implementations of FVA, such as VFFVA [81].

## Conclusion

In this work, we presented an algorithm that performs gap-filling while permitting metabolic interactions in microbial communities. Metabolites produced from one organism can either be released to the environment or used by another member of the community. Therefore, one of the key characteristics of the community gap-filling method is that it attempts to identify if interspecies exchanges of metabolites can compensate for insufficient metabolic capabilities of the community members, instead of just adding biochemical reactions from reference databases to the metabolic models of the community members. Our results showed that the algorithm can successfully predict both cooperative and competitive metabolic interactions in microbial communities and comply with experimental measurements and observations. With the ability to predict elusive metabolic interactions, the community gap-filling method can be used to generate hypotheses about potential metabolic relationships among community members. This direction is useful not only for the improvement of metabolic models, but also for understanding functions at the community level and providing valuable information for selective media composition that can enable the isolation of pure cultures for microorganisms.

The community gap-filling method can be further applied on communities with more than two members, even though such a scale-up could probably require a more computationally efficient reformulation. A way to use the community gap-filling method for the study of microbial communities could be the following. Applying the community gap-filling method for the simulation of a microbial community on different media can generate plenty of data on the function of the community in different environments. Evaluating these results to identify the reactions added to the models, as well as the metabolic interactions between the models, can offer an insight on which reactions are necessary for the metabolic reconstructions and which metabolic interdependencies are prevalent in the community. Such predictions can be further tested experimentally and can be used for improving gene-annotations by identifying gene functions, not based on sequence similarities, but rather on a growth restoration strategy.

## Supporting information

**S1 Appendix. Creation of glucose and acetate utilizer strains from the *E. coli* core model**.

**S2 Appendix. Reconstruction and individual curation of the metabolic models of *Dehalobacter* sp. CF and *Bacteroidales* sp. CF50**.

**S1 File. core E**.**coli community solutions**.**mat**. Alternative solutions for the core *E. coli* community.

**S2 File. gut-microbiome community solutions**.**mat**. Alternative solutions for the community of *B. adolescentis* ATCC 15703 and *F. prausnitzii* A2-165.

**S3 File. ACT-3 community solutions**.**mat**. Alternative solutions for the community of *Dehalobacter* sp. CF and *Bacteroidales* sp. CF50.

**S1 Table. Deleted and Exchange Reactions for *E. coli* glucose utilizer**.

**S2 Table. Deleted and Exchange Reactions for *E. coli* acetate utilizer**.

**S3 Table. Exchange Reactions for *B. adolescentis* ATCC 15703**.

**S4 Table. Exchange Reactions for *F. prausnitzii* A2-165**.

**S5 Table. Uptake Rates for ACT-3 community**.

**S6 Table. Constrained and Exchange Reactions for *Dehalobacter* sp. CF**.

**S7 Table. Exchange Reactions for *Bacteroidales* sp. CF50**.

**S8 Table. Reaction Fluxes from the best solution for the core *E. coli* community and Reactions Added to the models**.

**S9 Table. Fluxes of Exchange Reactions from the ten best solutions for the core *E. coli* community**.

**S10 Table. Fluxes of Added Database Reactions from the ten best solutions for the core *E. coli* community**.

**S11 Table. Reaction Fluxes from the best solution for the community of *B. adolescentis* ATCC 15703 and *F. prausnitzii* A2-165 and Reactions Added to the models**.

**S12 Table. Fluxes of Exchange Reactions from the ten best solutions for the community of *B. adolescentis* ATCC 15703 and *F. prausnitzii* A2-165**.

**S13 Table. Fluxes of Added Database Reactions from the ten best solutions for the community of *B. adolescentis* ATCC 15703 and *F. prausnitzii* A2-165**.

**S14 Table. Reaction Fluxes from the best solution for the community of *Dehalobacter* sp. CF and *Bacteroidales* sp. CF50 and Reactions Added to the models**.

**S15 Table. Fluxes of Exchange Reactions from the ten best solutions for the community of *Dehalobacter* sp. CF and *Bacteroidales* sp. CF50**.

**S16 Table. Fluxes of Added Database Reactions from the ten best solutions for the community of *Dehalobacter* sp. CF and *Bacteroidales* sp. CF50**.

**S1 Fig.**
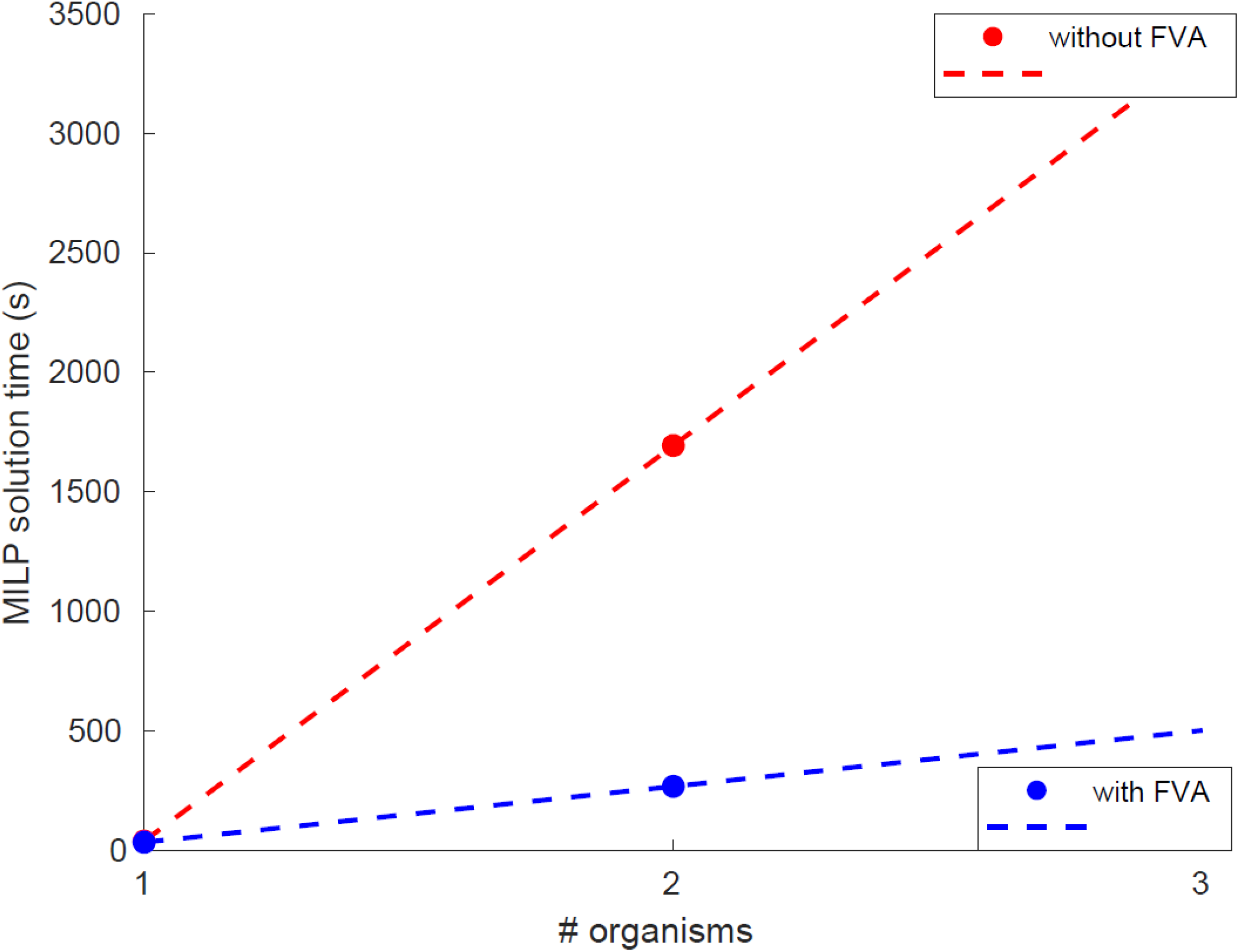
Time needed for the solution of the MILP problem with and without the prior use of FVA for the reduction of the solution space, depending on the number of organisms in the community. As the number oforganisms that make up the microbial community increases, the community gap-filling algorithm needs increasingly more time to solve the MILP problem that restores growth in the community. However, the increase in solving time is significantly reduced with the use of our community gap-filling method that reduces the solution space of the MILP problem by performing FVA for each organism compartment of the community before formulating the optimization problem. (The presented measurements were performed with the models from the toy *E. coli* community.)

**S2 Fig.**
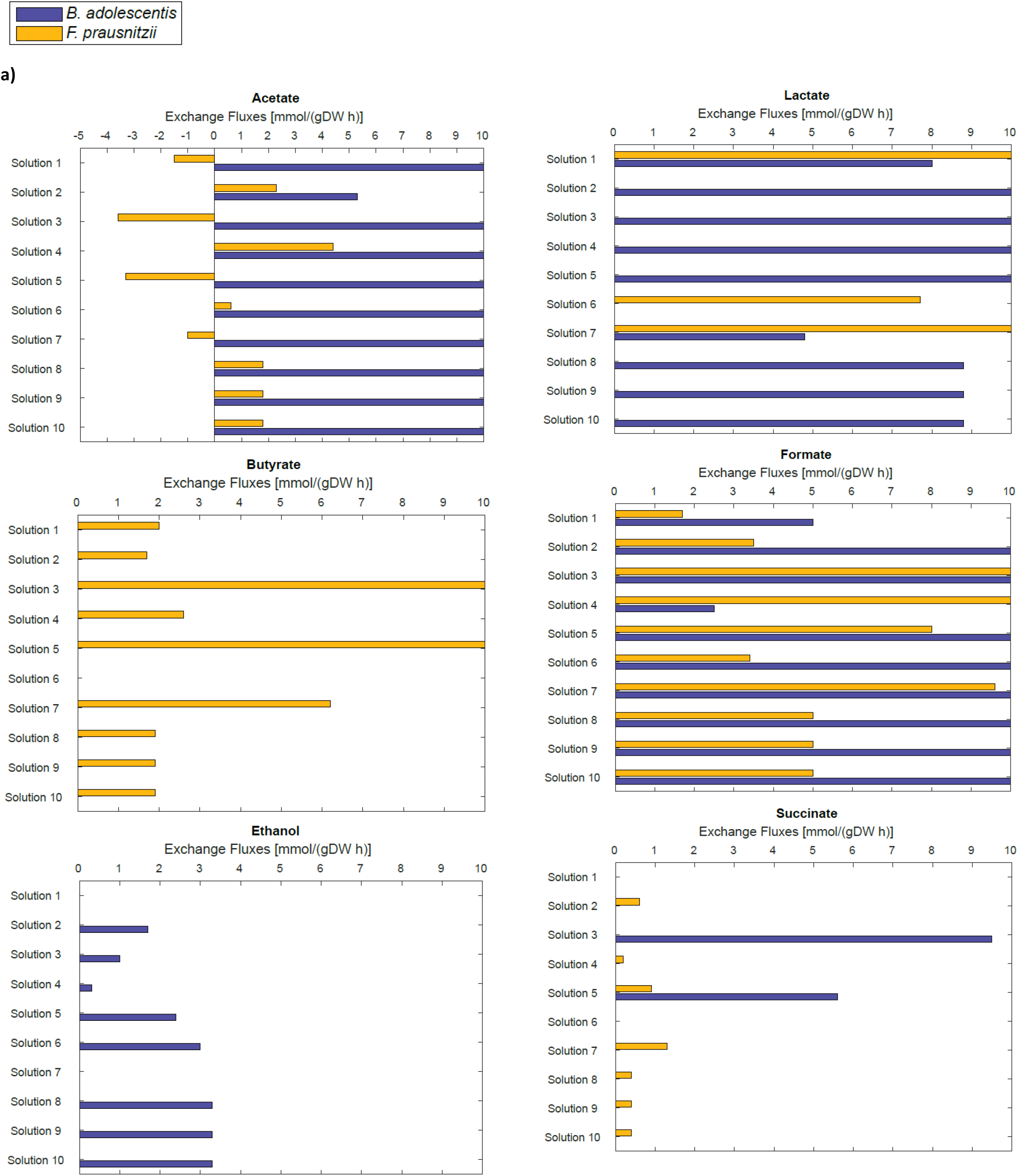
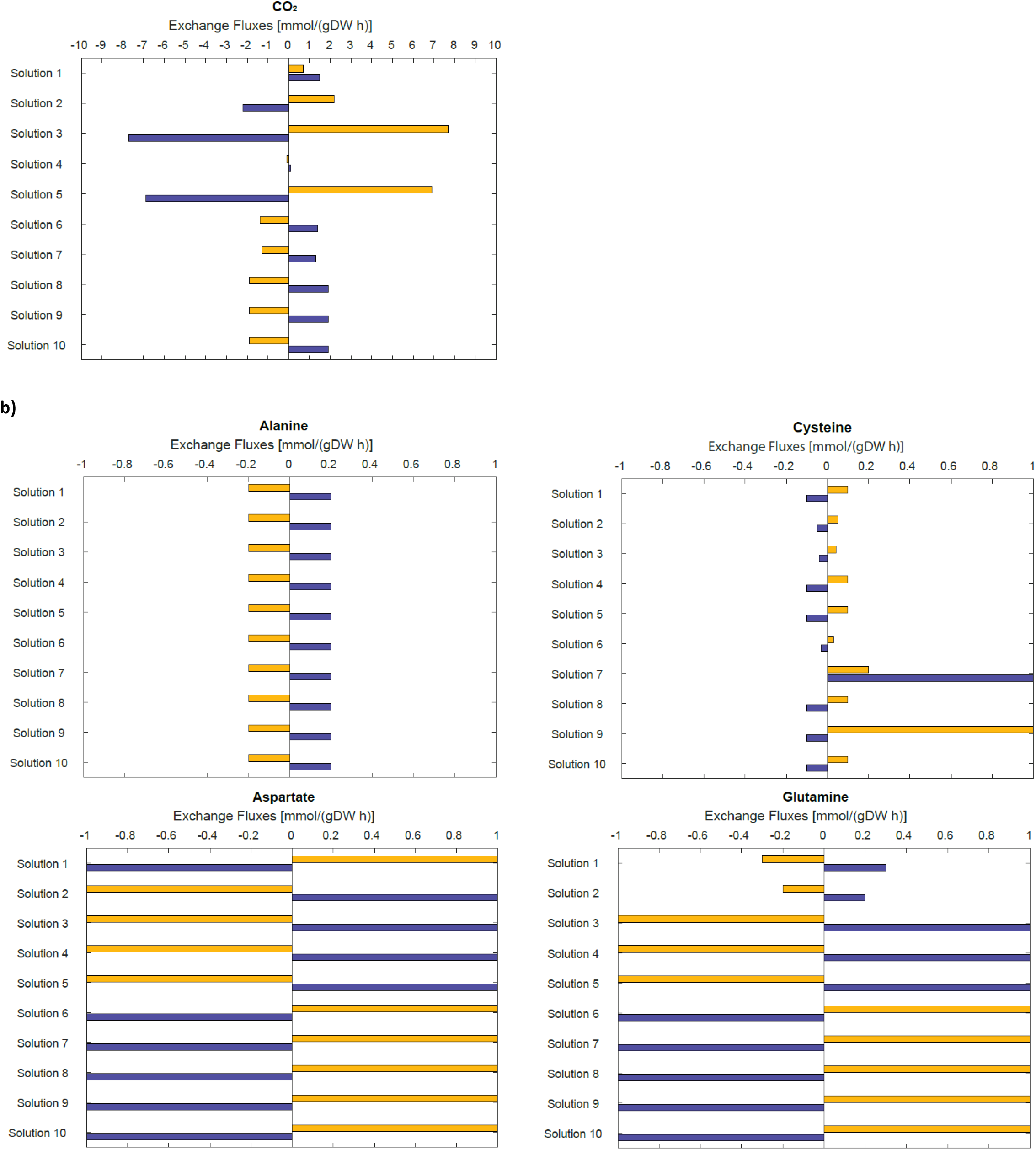
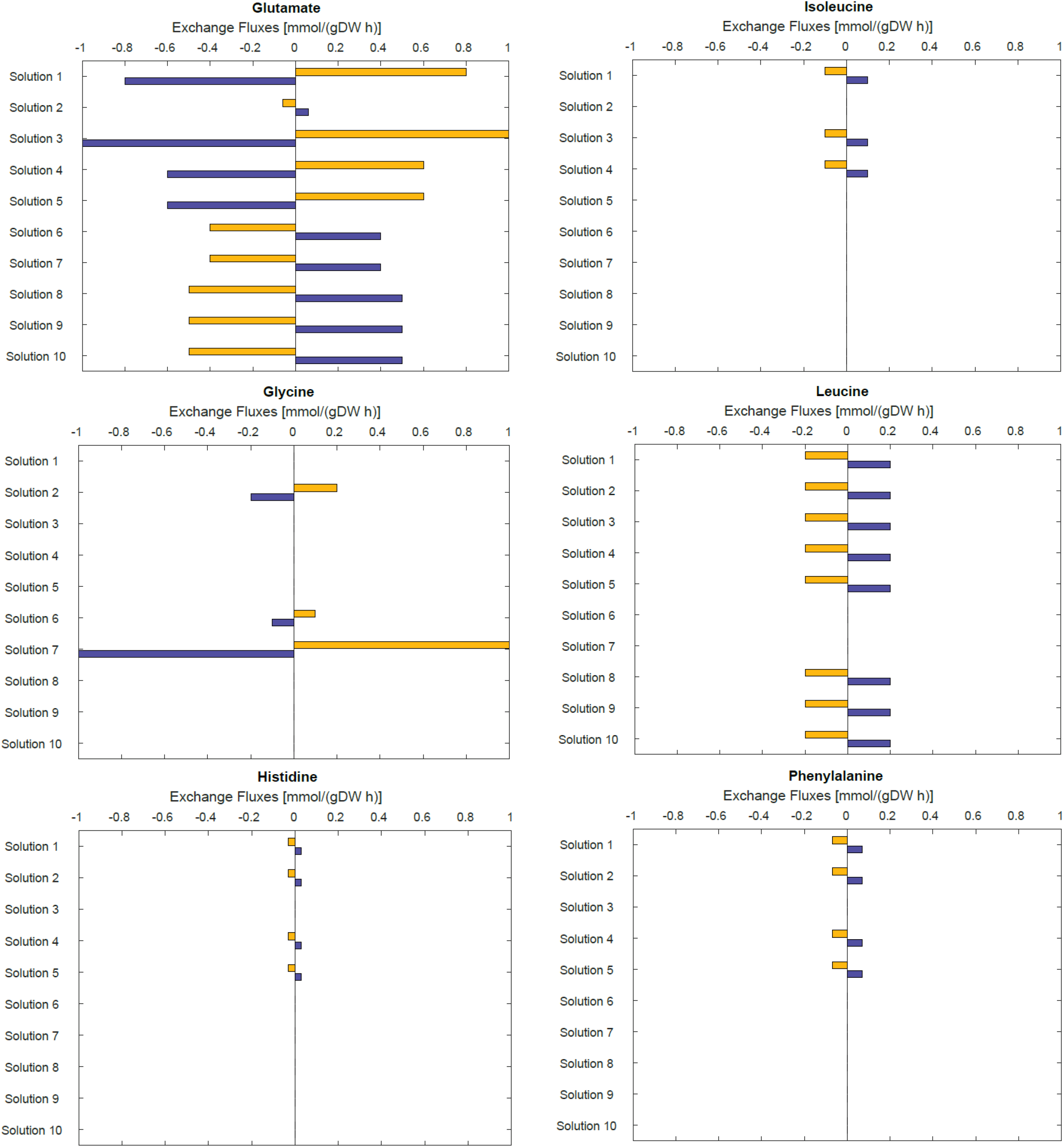
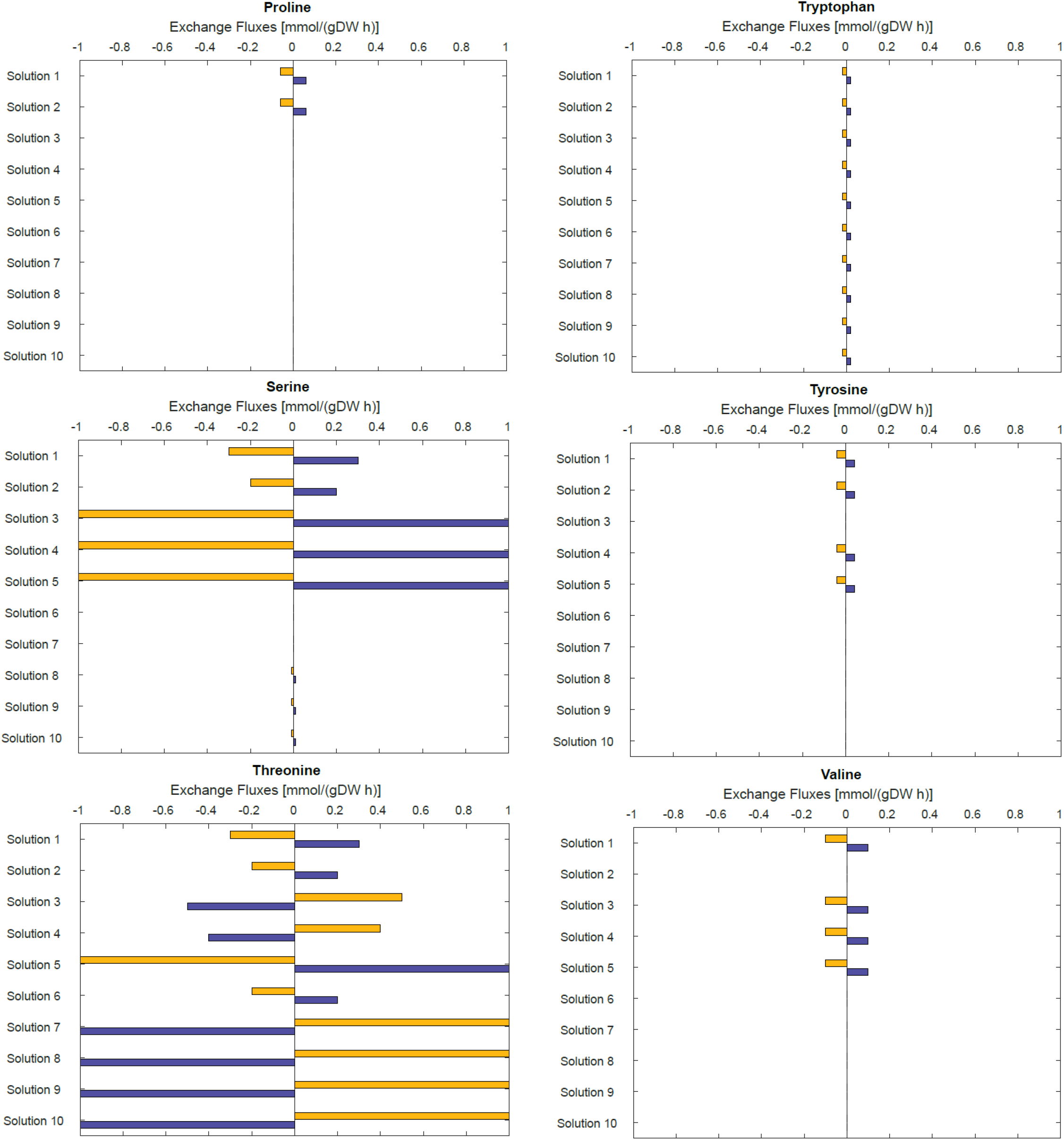
Fluxes of Exchange Reactions from the ten best solutions for the community of *B. adolescentis* ATCC 15703 and *F. prausnitzii* A2-165. **a)** Fluxes of exchange reactions for the SCFAs acetate, butyrate, ethanol, lactate, formate, and succinate, and CO_2_. **b)** Fluxes of exchange reactions for the amino acids alanine, aspartate, cysteine, glutamine, glutamate, glycine, histidine, isoleucine, leucine, phenylalanine, proline, serine, threonine, tryptophan, tyrosine, and valine.

**S3 Fig.**
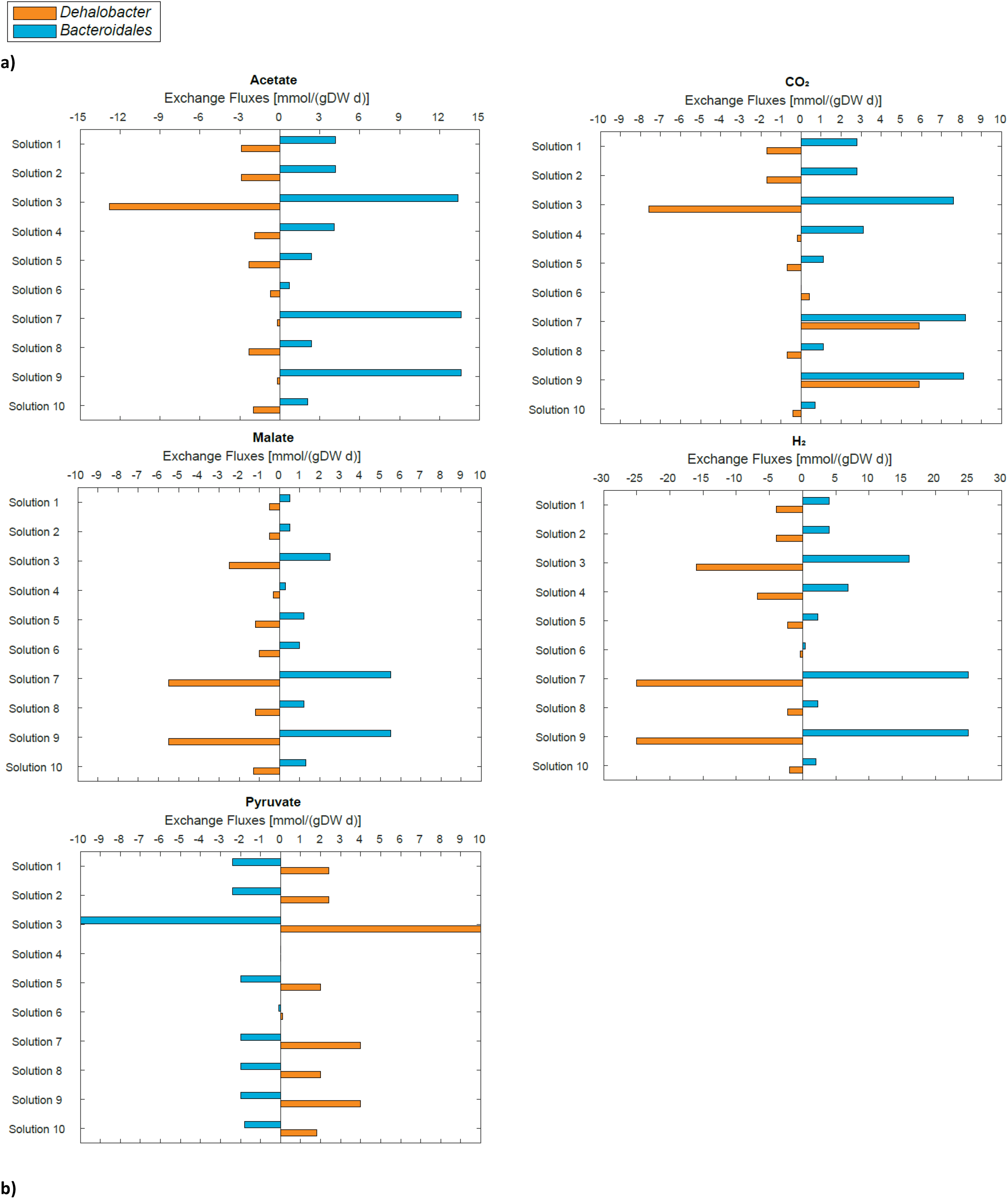
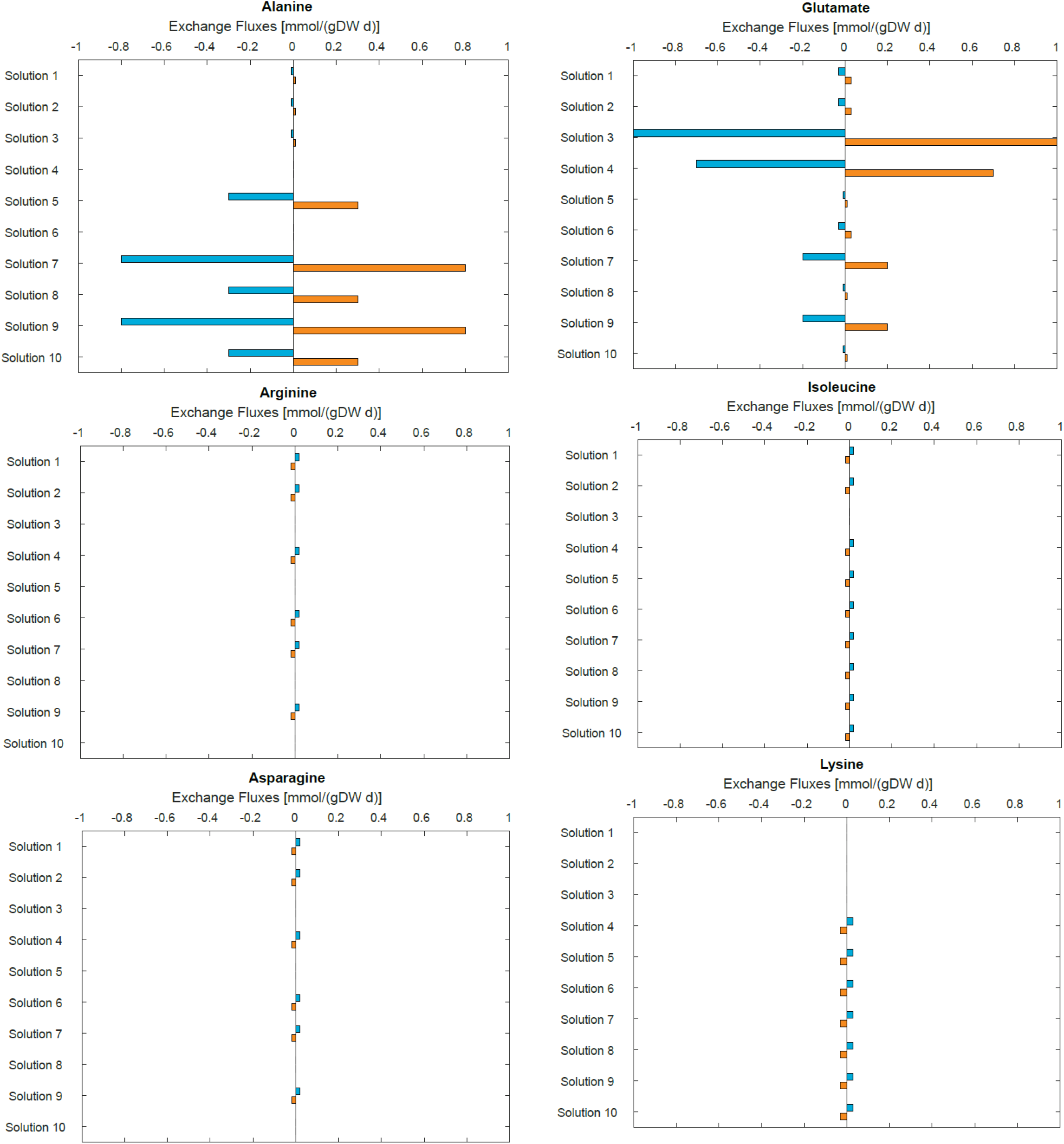
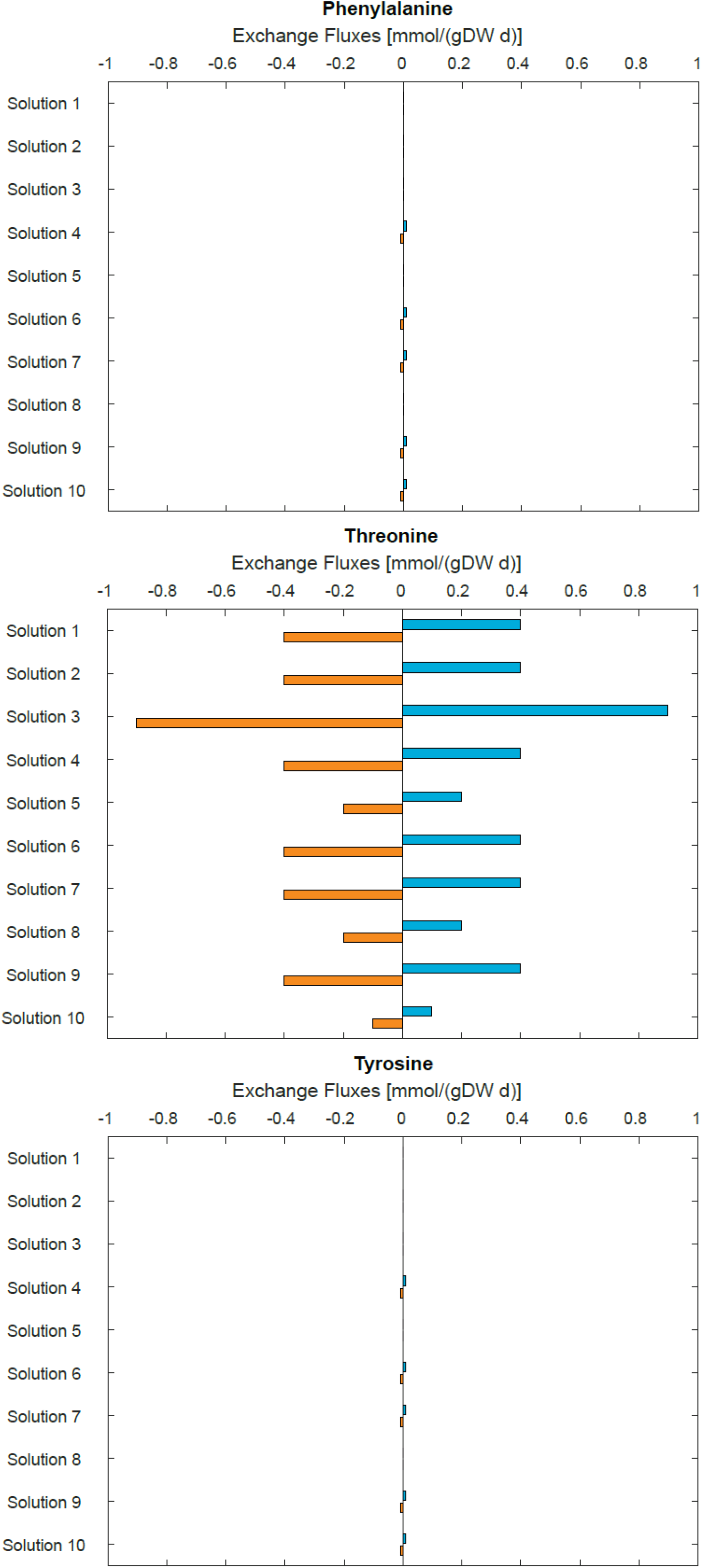
Fluxes of Exchange Reactions from the ten best solutions for the community of *Dehalobacter* sp. CF and *Bacteroidales* sp. CF50. **a)** Fluxes of exchange reactions for the organic acids acetate, malate, and pyruvate, CO_2_ and H_2_. **b)** Fluxes of exchange reactions for the amino acids alanine, arginine, asparagine, glutamate, isoleucine, lysine, phenylalanine, threonine, and tyrosine.

## Availability

The code used to perform this study can be found on GitHub: https://github.com/LMSE/Community_Gap-Filling.

## Acknowledgments

This work was funded by Natural Sciences and Engineering Research Council of Canada (NSERC), Genome Canada, and Ontario Ministry of Research and Innovation. We thank Kevin Correia, Kaushik Raj Venkatesan, and Anna Khusnutdinova for fruitful conversations and editing.

